# Effective normalization for copy number variation in Hi-C data

**DOI:** 10.1101/167031

**Authors:** N. Servant, N. Varoquaux, E. Heard, JP. Vert, E. Barillot

**Affiliations:** Institut Curie, Paris, France; INSERM, U900, Paris, France; Mines ParisTech, PSL-Research University, CBIO-Centre for Computational Biology, Fontainebleau, France; Department of Statistics, University of California, Berkeley, USA; Berkeley Institute for Data Science, Berkeley, USA; CNRS UMR3215, Paris, France; INSERM U934, Paris, France; Ecole Normale Superieure, Department of Mathematics and Applications, Paris, France

## Abstract

Normalization is essential to ensure accurate analysis and proper interpretation of sequencing data. Chromosome conformation data, such as Hi-C, is not different. The most widely used type of normalization of Hi-C data casts estimations of unwanted effects as a matrix balancing problem, relying on the assumption that all genomic regions interact as much as any other. Here, we show that these approaches, while very effective on fully haploid or diploid genome, fail to correct for unwanted effects in the presence of copy number variations. We propose a simple extension to matrix balancing methods that properly models the copy-number variation effects. Our approach can either retain the copy-number variation effects or remove it. We show that this leads to better downstream analysis of the three-dimensional organization of rearranged genome.

## Background

The spatial organization of the genome and the physical interactions occurring within and between chromosomes are known to play an important role in gene regulation and in genome function in general. The organization and the folding of mammalian chromosomes within the nucleus involve multiple hierarchical chromatin structures (see Bonev and Cavalli [2016] for a review). At the megabase-scale, chromosomes are divided to genomic compartments of active and inactive chromatin, respectively associated with gene-rich, actively transcribed regions and gene-poor, silent regions [Lieberman-Aiden et al., 2009, Rao et al., 2014]. The arrangement of these compartments varies across physiological conditions and cell differentiation [Dixon et al., 2015, Barutcu et al., 2015]. At the sub-megabase scale, chromosomes are partitioned into topological associated domains (TADs). These functional units of regulation are well conserved both across cell types and between mammals [Nora et al., 2012, Dixon et al., 2012, Rao et al., 2014, Dixon et al., 2015]. TAD boundaries are frequently associated with the presence of the CTCF binding factor, itself also involved in the establishment of chromatin loops between convergent target sites [Rao et al., 2014]. These chromatin loops are also commonly associated with promoter-enhancer contacts and therefore with gene activation (see Bouwman and de Laat [2015] for a review).

Given the important recent insights that chromosome conformation techniques have provided into 3D genome organization in a normal context, the application of such approaches to a disease context offers great promises to explore the effect of perturbations in 3D genomic organization on cell regulation (see Krijger and de Laat [2016] for a review). At a high enough resolution, such techniques can be used to characterize links between disease-associated sequence variants and the gene regulatory landscape. For example, structural variants can disrupt the boundaries between TADs, and consequently act as driver events in the mis-regulation of associated gene expression [Franke et al., 2016, Lupiáñez et al., 2016].

Over the past decade, both major advances in high-throughput sequencing techniques and the availability of data from large patient cohorts across multiple cancer types have enabled a comprehensive and systematic exploration of genomic and epigenomic landscapes of a wide variety of cancers. While cancer has been shown to have a genetic component, our appreciation of the inherent epigenetic complexity is more recent and has dramatically increased over the last few years. At the genetic level, cancer is frequently associated with the sequential acquisition of somatic variants, both at single nucleotide and at the copy number levels [Ciriello et al., 2013]. The different alterations that characterize tumors are usually caused by a few functional driver events, which occur among many non-functional passenger events, mainly located in the non-coding part of the genome [Vogelstein et al., 2013]. One of the exciting discoveries that has emerged from systematic sequencing of cancer genomes was the high frequency of mutations in genes known to regulate epigenetic processes such as chromatin associated proteins, DNA methylation, or histone variants and modifications [Plass et al., 2013]. The contribution of altered epigenomes in the process of tumorigenesis is thus at last being unraveled thanks to the combination of genomic and epigenomic interrogation. More recently, genetic and epigenetic alterations in the non-coding part of the genome, including distal regulatory elements such as enhancers or insulators, have been reported and found to impact gene expression in cancer [Taberlay et al., 2014]. This has led to intense interest in the spatial proximity and 3D organization of cancer genomes. Losada [2014] reviews the effect of somatic mutations in cohesin complex proteins (which play a citical role in TADs organization and chromosome looping) in various types of cancer. Gröschel et al. [2014] and Taberlay et al. [2016] describe how disruptions in genome organization (respectively in leukemia and prostate cancer) lead to major epigenetic and transcriptional changes. Lastly, Hnisz et al. [2016], Weischenfeldt et al. [2017], Beroukhim et al. [2016] show how disruptions in long range DNA looping and genome rearrangements lead to enhancer hijacking and Flavahan et al. [2016] link insulator dysfunctions to oncogene activation in cancer. Thus, changes in chromosome conformation at different scales are now considered as key players in cancer, as well as important potential biomarkers.

Developing accurate and quantitative methods to analyze the chromatin conformation derived from disease-associated cells/tissues is therefore of increasing interest to a wide community of researchers and pathologists. In addition to standard microscopy approaches, several 3C-based methods have been proposed: these rely on digestion and religation of fixed chromatin to estimate the probability of contact between two genomic loci (see Ramani et al. [2016] for a review). In Hi-C experiments, the contact frequencies between two genomic loci are roughly proportional to the reads counts observed between two regions after sequencing [Lieberman-Aiden et al., 2009]. However, as is the case for many high-throughput technologies, the raw contact frequencies are affected by technical biases such as GC content, mappability, or restriction fragment size [Yaffe and Tanay, 2011]. Estimating and correcting these biases is therefore an important step in ensuring accurate downstream analysis. In the past few years, several methods and packages have been developed to normalize Hi-C data (see Ay and Noble [2015] for a review). These methods fall into two main categories: explicit factor correction methods or matrix balancing algorithms (such as the iterative correction (ICE) method [Imakaev et al., 2012]). In the context of cancer Hi-C data, an additional perturbation related to chromosomal rearrangements must be considered. Amplified genomic regions have a greater chance of being pull-down during the library preparation, while genomic regions with lower copy numbers are more difficult to detect. To date, such copy number variants (CNVs) are usually ignored in cancer Hi-C data normalization, although they raise interesting and important questions both at the biological and methodological levels. The real impact of CNVs on contact frequencies remains difficult to assess. For instance, a tandem amplification does have a very different impact on local chromatin organization compared to the gain of a complete chromosome. Similarly, a genomic duplication could lead to different changes in contact frequencies depending on whether the event occurs within a TADs or across/at its boundary [Franke et al., 2016]. Addressing the question of CNVs during normalization is therefore an important challenge in the analysis of Hi-C data and its interpretation in the context of genetic and epigenetic misregulation in disease.

The question of how copy number signal should be treated mainly depends on the related biological questions. The first strategy is to consider the copy number effect as an unwanted effect, and to remove it during the normalization step [Wu and Michor, 2016]. This strategy indeed makes sense for the detection of a genomewide list of significant contacts, or for the direct comparison of samples with different chromosomal rearrangement profiles. On the other hand, signal from copy number alterations can also be considered as an important biological information, that can be of interest for 3D modeling, genome reconstruction of cancer cells, or to simply further characterize the genomic landscape of a tumor [Harewood et al., 2017].

Here, we propose to further explore the impact of CNVs on Hi-C data and provide tools that deal with its effect on data normalization. First, we develop a model simulating large copy number rearrangements on a diploid Hi-C contact map. Using such simulated data, we demonstrate that the naive matrix balancing algorithm which is commonly used to normalize Hi-C data, cannot be applied to cancer Hi-C data. We then propose two methods that extend the ICE algorithm and correct the data from systematic biases, either considering the CNVs as a bias to remove or as an interesting signal to conserve in the data structure. Finally, we apply these methods to several disease associated Hi-C data sets, demonstrating their relevance.

## Results

### Simulating the effect of copy number variations on Hi-C data

Due to the large number of genomic and epigenomic factors possibly involved, predicting the true effect of copy-number variations on the 3D organization of the genome is challenging. We propose a simple mathematical model to simulate the effect of abnormal karyotypes on a diploid Hi-C data set by estimating the enrichment in interaction due to copy number variation (see Methods and Figure 1).

**Figure 1:**
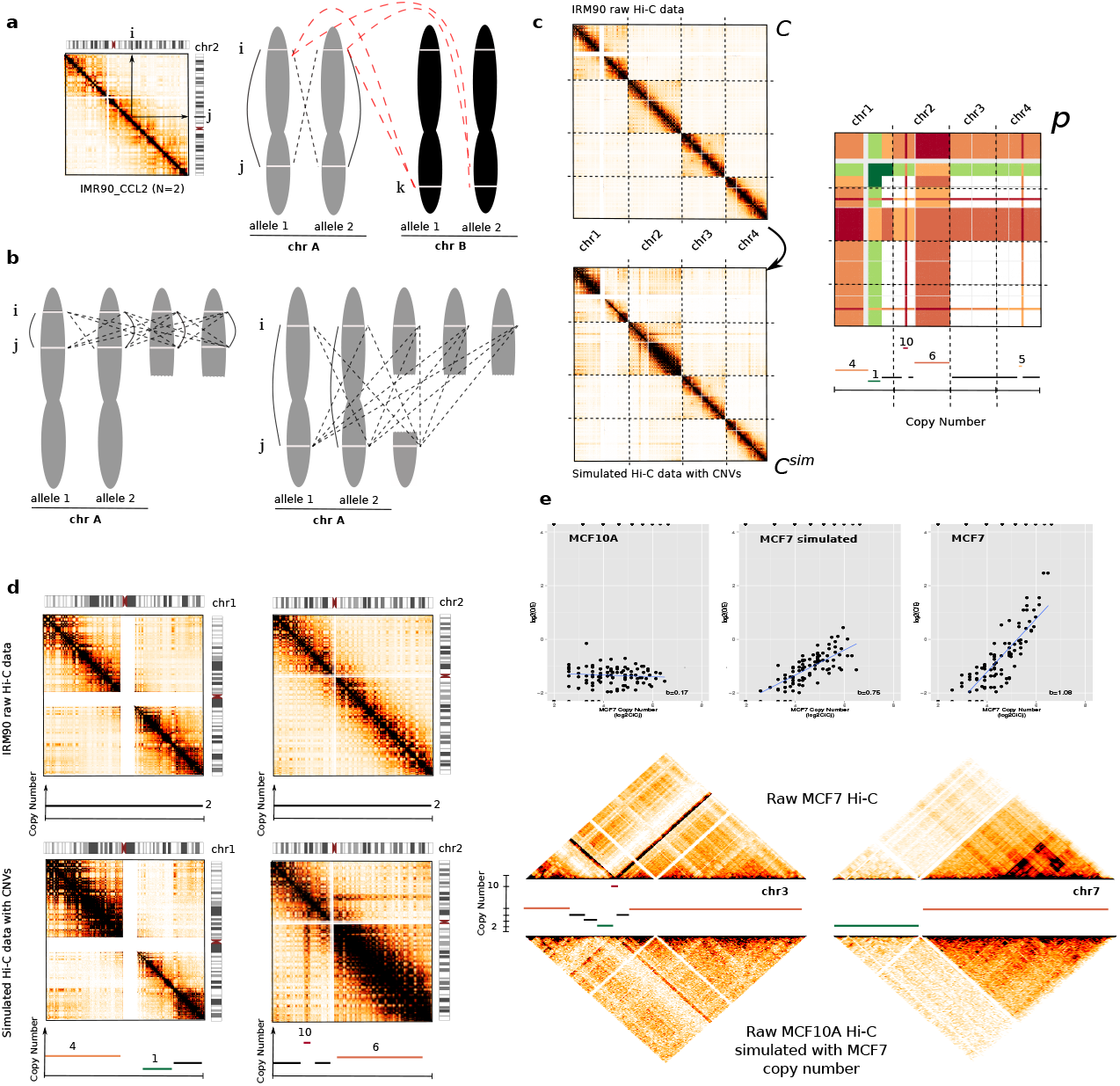
Simulation of cancer Hi-C data. **a.** In diploid Hi-C data, the contact frequency measured between two loci *i* and *j* is equal to the sum of 2 *cis* interactions (black solid lines) occurring within an individual allele and of 2 *trans* interactions between homologous chromosomes (*transH*, black dashed lines). In addition, the contact frequency observed in *trans* between loci *i* and *k* is the sum of 4 interactions between non homologous chromosomes (red dashed lines). **b.** In the context of segmental rearrangement, these properties can be extended and generalized if loci *i* and *j* belong to the same DNA segment, or to different segments (see Methods and Figure S1). **c.** Simulation of cancer Hi-C data from normal diploid (C) data by calculating the scaling factor matrix (p). Colors in scaling factor matrix represents the level of gains (red) and loss (green) to simulate. For each interaction 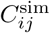, the simulated count is finally estimated using a binomial downsampling method of probability (see Methods). **d.** Intra-chromosomal maps of chromosome 1 and 2 before (top) and after (bottom) simulation of copy number changes. Copy number effects are characterized by blocks of high/lower signal. Overall, the simulation conserves the structure and the counts/distance properties of the Hi-C maps. **e.** Validation of the simulation model using Hi-C data from MCF10A cell line from which we simulated the expected copy number of MCF7 cancer cell line. We observed a positive correlation between the raw log2 O/E (Observed/Expected) ratios and log2 multiplicative copy number in 1 Mb resolution Hi-C maps, on both simulated and real MCF7 Hi-C data. Looking at the intra-chromosomal maps of chromosomes 3 and 8 demonstrates that our model efficiently simulates large copy number events.

In order to validate our simulation model, we leverage available Hi-C data from two epithelial cell lines: the MCF7 breast cancer cell line and the MCF10A nearly diploid, non-tumorigenic cell line [Barutcu et al., 2015]. We extract MCF7’s copy number information from Affymetrix SNP6.0 array, filtering out any altered segments lower than the MCF10A’s Hi-C map resolution (1 Mb) and apply our simulation model on the normal-like data, thus obtaining a simulation of MCF7’s abnormal karyotype. We then compare our simulated results with the real MCF7 Hi-C data set. As expected, the contact counts (for both the simulated data and the real data) are correlated with the copy number (Figure 1e). Both our simulations and the real data show blocks of higher/lower contact frequencies in regions affected by large copy number variants. In average, the intra-chromosomal maps of simulated and real MCF7 data have a Spearman correlation of 0.70 (0.54-0.84). We then summarize both data in 1D by summing the contact frequencies over each row. Overall, the simulated MCF7 profile is well correlated with the profile of real MCF7 Hi-C data (Spearman cor=0.88, Figure S3). We can observe that the sum of interactions for a genomic window is proportional to the copy number. This is expected, as a genomic region in multiple copies have a greater chance of being seen interacting with another region. Interestingly, for highest copy number, both profiles increase concurrently, but not at the same rate (Figure 1e). One explanation would be that these regions of very high copy number correspond to tandem focal amplifications. Their linear proximity on the genome would therefore explain the massive increase of contact frequencies that we observed in real data, and which are not modeled by our simulation.

Overall, those observations leads us to believe our simulation method appropriately models the effect of copy number variations on Hi-C data.

### The ICE normalization is not suitable for cancer Hi-C data

We then apply this simulation model to assess the ability of the ICE normalization method to correct for copy number variations. We simulate two data sets with different properties from the publicly available Human IMR90 Hi-C data [Rao et al., 2014]. First, we generate a highly rearranged data set, with segmental gains, losses and a focal amplification up to 10 copies (Figure 1c). Then, we simulate a case of aneuploidy with gain or loss of entire chromosomes (Figure S2a). While the simulation is performed genome-wide, we restrict the CNVs to the first chromosomes to ease the results interpretation and visualization.

Several methods have been proposed to remove unwanted technical and biological variations from Hi-C data, none of which are adequate for abnormal karyotype data. These methods fall into two groups. The first explicitly models sources of biases, and cast the normalization procedure as a regression problem [Yaffe and Tanay, 2011, Hu et al., 2012]. The second group leverages a small number of hypothesis on the bias and on the properties of Hi-C data to formulate the normalization procedure as matrix-balancing problems: these do not assume any specific sources of biases, and are (as long as the hypothesis are fulfilled) able to correct for any factors affecting contact frequencies [Imakaev et al., 2012, Cournac et al., 2012]. Among those methods, the iterative correction method (ICE, Imakaev et al. [2012]) has been successfully applied to many diploid Hi-C data sets. ICE relies on two assumptions: (1) the bias between two regions *i* and *j* can be represented as the product of individual biases of these regions: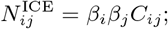 (2) each bin should interact approximately the same number of times: 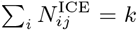 where *C* represents the raw count matrix, *N* ^ICE^ the ICE normalized count matrix, *β* the bias vectors and *k* a constant.

In this section, we explore the effects of ICE on Hi-C data in the presence of copy number variations. To do this, we leverage the simulated data set previously described (Figure 1c, Supp. Figure S2a) for which the ground-truth normalized data is found by applying ICE to the original diploid data. We are thus able to assess the performance of ICE to correct for unwanted sources of variation, including the copy number, by comparing the obtained matrices to the ground-truth.

Before normalizing the data with ICE, we first represent the data in 1D, by summing each row of the matrix. As previously mentionned, the sum of genome-wide interactions per bin is proportional to the copy number (Figure 2a). After applying ICE, each genomic region now interacts the same number of times genome-wide, as expected. However, ICE leads to an imbalance between *cis* and *trans* contact counts. As shown on Figure 2b, we observe that the *cis* contact counts are now depleted for regions with high copy number, and *trans* contact counts are enriched. On the other hand, lost regions now present higher contact probabilities than gain regions in *cis*. The same conclusions can be made in the context of aneuploidy (Figure S2). However, we notice that in this case, ICE could yield to the expected results if the analysis is restricted to intra-chromosomal contacts. We thus conclude that ICE is not adapted to correct for segmental copy number effect, and that, more importantly, it can lead to a misinterpretation of the contact probability between rearranged regions.

**Figure 2:**
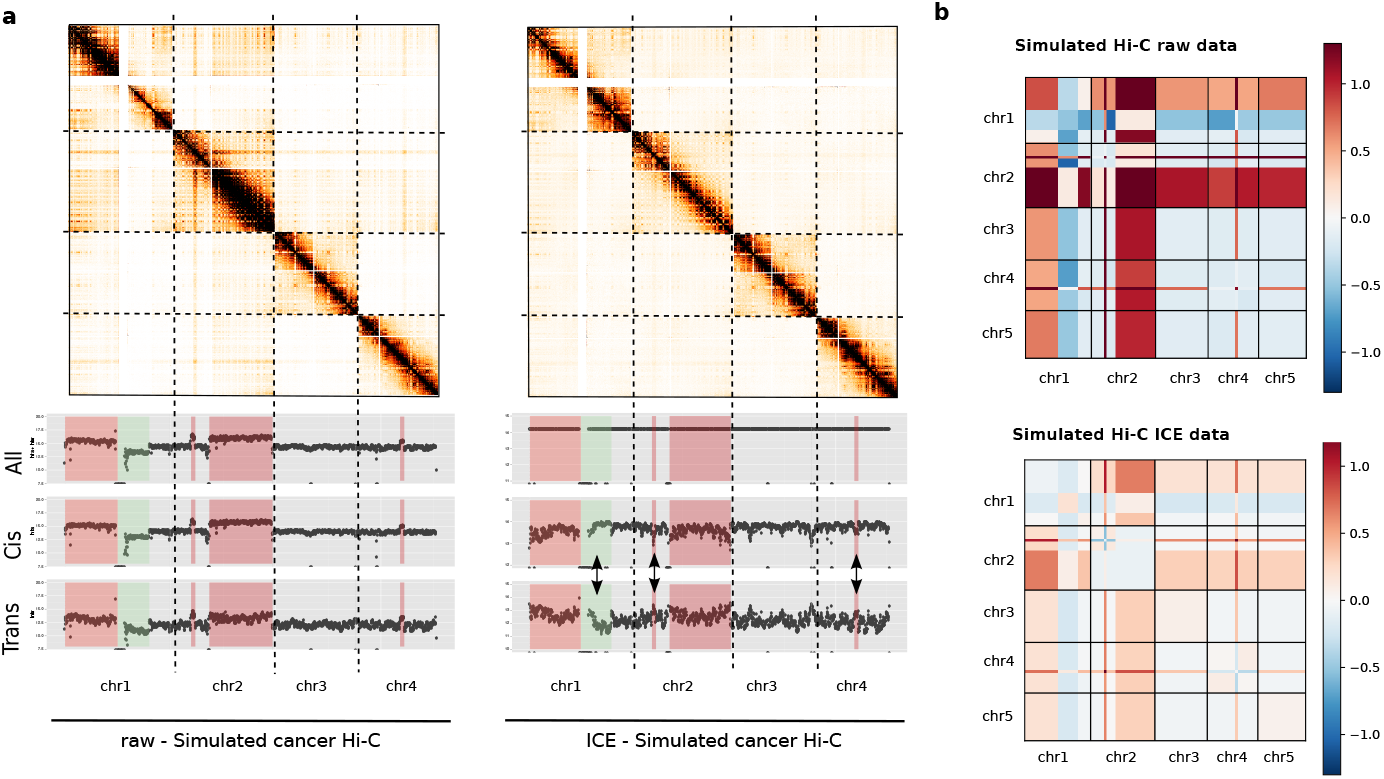
Impact of matrix balancing normalization on simulated cancer Hi-C data. **a**. Simulated Hi-C contact maps (500 kb resolution) of the first four chromosomes and contact frequencies presented as the sum of genome-wide contacts per locus, using either all (inter and intra-chromosomal), *cis* (intra-chromosomal) or *trans* (inter-chromosomal) contacts. Rearranged regions are highlighted in red (gain) or green (loss). The 1D profile of ICE data is constant genome-wide as expected under the assumption of equal visibility. However, the iterative correction on simulated cancer data results in an shift of contacts between altered regions (arrows). **b**. Block-average error matrix of simulated raw and ICE cancer data (150 Kb resolution) (See supp method 1.4). The iterative correction does not allow to correct for segmental copy number bias.

If the downstream analysis is restricted to intra-chromosomal interactions, one may ask whether applying ICE independently to each intra-chromosomal maps could mitigate the introduction of biases. We therefore independantly normalized by ICE all intra-chromosomal maps. However, although the effects are less strong, we observe the same phenomenon in complex rearrangements (Figure S4).

Altogether, these results demonstrate that ICE does not properly normalize data with abnormal karyotype and that its use is therefore not recommended in the context of cancer Hi-C data.

### LOIC: a novel normalization strategy for cancer Hi-C data

As discussed above, ICE relies on the assumption of equal visibility of each genomic bin. In the presence of copy number variations, this assumption does not hold: genomic bins with higher copy number variations will interact overall more than genomic bins of lower CNVs. In addition, the copy number effect between loci *i* and *j* (*B_ij_*), cannot be decomposed as the product of an effect in loci *i* and an effect in loci *j*, therefy also violating the ICE hypothesis. Instead, we propose to extend the ICE model, saying that the assumption of equal visibility remains true across regions of identical copy number. In addition, biases associated to fragments (such as fragment length, GC-content, or mappability) are still decomposable into the product of two region specific biases.

We thus first propose to extend ICE by assuming that the sum of contacts for a given genomic bin is constant across genomic bins of identical copy number (Figure 3a) 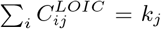where *k_j_* is the interaction profile associated to the copy number of *j* (see Methods). We refer to this method as a local iterative correction (LOIC). When there is no copy number aberration, LOIC solves exactly the same problem as ICE.

**Figure 3:**
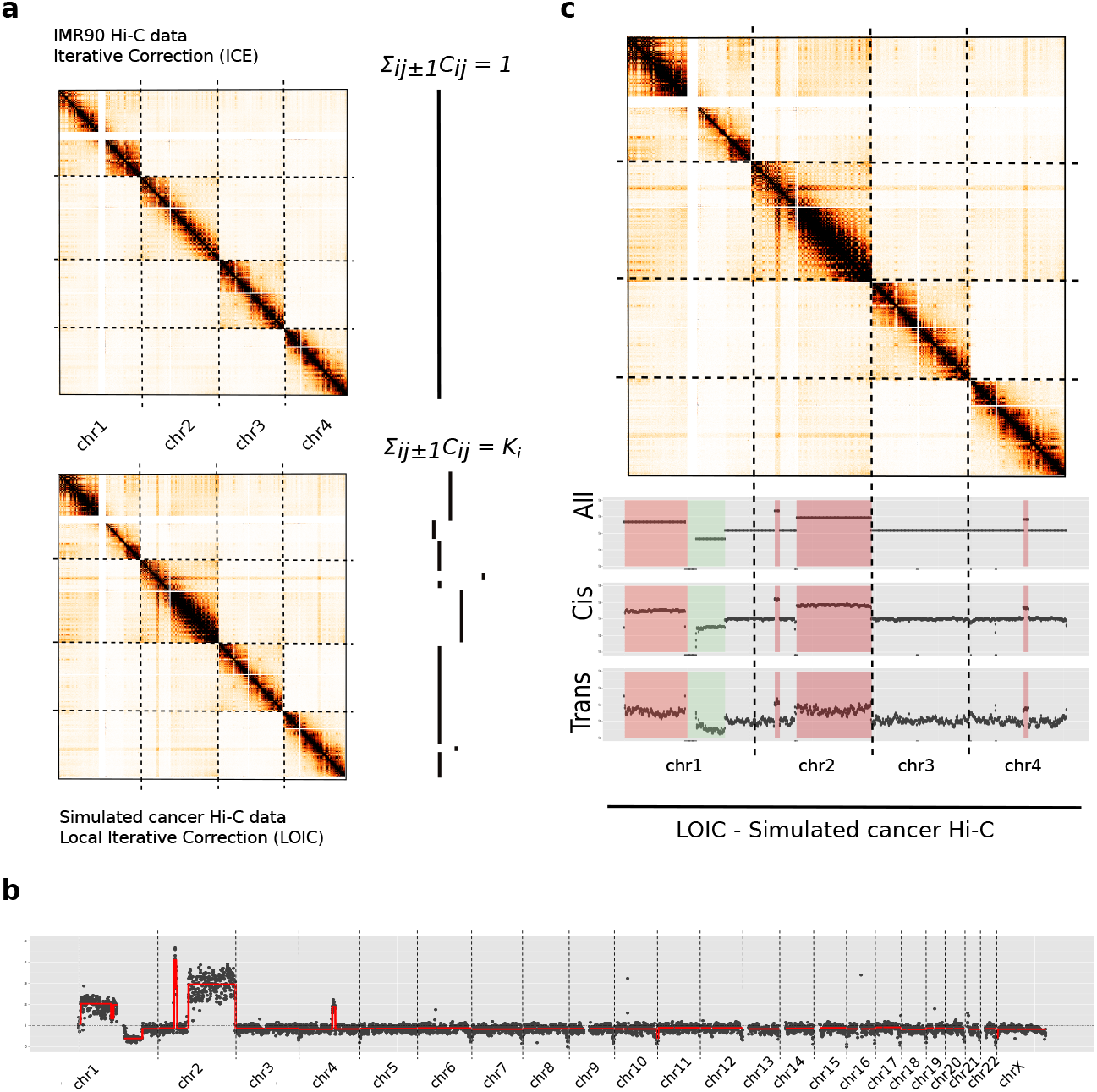
Generalization of matrix balancing algorithms for cancer Hi-C data. **a.** Rationale of LOIC method versus standard ICE method. The LOIC method extends the ICE normalization by constraining the genome-wide Hi-C 1D profile to follow the copy number signal. **b.** Segmentation of the Hi-C 1D genomewide profile of simulated cancer data. The red line represents the smoothing line that estimate the copy number level. **c.** LOIC normalized Hi-C contact maps of simulated data on the first four chromosomes. The 1D profiles are represented by the sum of genome-wide contacts at each locus using either all (inter and intrachromosomal), *cis* (intra-chromosomal) or *trans* (inter-chromosomal) contacts. As a results, we can see that the LOIC method allows to normalize cancer Hi-C data keeping into account the copy number information.

The LOIC procedure requires to identify DNA segments of equal copy number. External sources of data (such as genome sequencing or microarray data) can be used to infer DNA breakpoints along the genome, and thus to define the DNA segments of equal copy number. If such data is not available, we propose to directly infer the copy number from the Hi-C data (see Methods). Applying our method to the simulated data yields a copy number estimation well correlated with the profile used for simulation (Spearman cor=0.64) (Figure 3b). To further validate this step, we apply the segmentation procedure to the IMR90 diploid data set. We obtain a nearly uniform copy number profile (Figure S6c). Interestingly, we frequently observe a decrease of contacts at telomeric regions which can therefore leads to a breakpoint in the segmentation. This telomeric pattern is expected, even in a diploid sample, as the assumption of equal visibility in these regions can be discussed.

We then apply the LOIC procedure to our highly rearranged simulated Hi-C data set using the breakpoint positions estimated by our segmentation procedure (Figure 3c). As expected, we observe that the genome-wide sum of contacts of each bin is proportional to the copy number, and that bins within a DNA segment are normalized to the same level of interactions. The previous effects observed on *cis* and *trans* sum of contacts with the standard ICE strategy no longer hold true. In addition, we calculate the effective fragment length, the GC content and the mappability features for each 500 Kb bin as already proposed [Hu et al., 2012], and then represent the average contact frequencies among those genomic features. Despite a few local enrichment due to CNVs, we observe that the LOIC normalization is as effective as the ICE normalization to correct for GC content, effective fragment length and mappability (Figure S5). We then turn to the aneuploid simulated data set. In the latter case, the LOIC allows to conserve the inter-chromosomal scaling factor due to CNVs. Differences in intra-chromosomal maps between ICE and LOIC remains negligible in this case and are related to the segmentation profile (Figure S6a,b).

To conclude, the LOIC strategy can be seen as a generalization of the ICE method. In that sense, applying both methods to a diploid data set leads to identical results.

### CAIC: estimating and removing the copy-number effect on cancer Hi-C data

In addition, we also propose to estimate and to correct the effect introduced by copy number changes. We assume that the copy number effect can be represented as a block constant matrix where each block is delimited by a copy number change (see Methods). In addition, we assume that, on average, each pair of loci interacts the same way as any pair of loci at the same genomic distance *s*. In summary, the raw interaction count *C_ij_* is roughly equal to the product of a CNV bias *B_ij_* and the expected contact count at genomic distance *s*: *C_ij_* ≃ *B_ij_ e_s_*_(*i,j*)_. We thus cast an optimization problem to find the CNV block biases *B* and the expected contact count at genomic distance *s* (see Methods). We refer to this method as CNV-Adjusted Iterative Correction (CAIC). We apply the CAIC normalization to the two simulated data sets. Looking at the 1D signal of the CAIC normalized data using the *cis* and *trans* data validates that the method tends to remove the CNV effect (Figure 4a). In addition, the unbalanced effect that we previously observed with the ICE normalized data disappears. We then divide the normalized contact matrices by the expected count matrices, thus removing the structure due to genomic proximity. Taking the average per block, we observe that the expected CAIC matrices are much more uniform than the ICE normalized matrices (Figure S7 and S8). On the aneuploid simulated data set, it is worth noting that ICE and CAIC yield very close results. We then compare the normalized contact maps to the ground-truth by computing the error matrix as well as three additional error measures (see Figure 4b and Supp Methods 1.4). We observe that the copy number effect is well removed both on the aneuploid (*l*_1_ = 2.291 *×* 10^6^, *l*_2_ = 3.532 *×* 10^5^ and *l*_max_ = 0.185) and highly rearranged data set (*l*_1_ = 2.055 *×* 10^6^, *l*_2_ = 2.536 *×* 10^5^ and *l*_max_ = 0.360).

**Figure 4:**
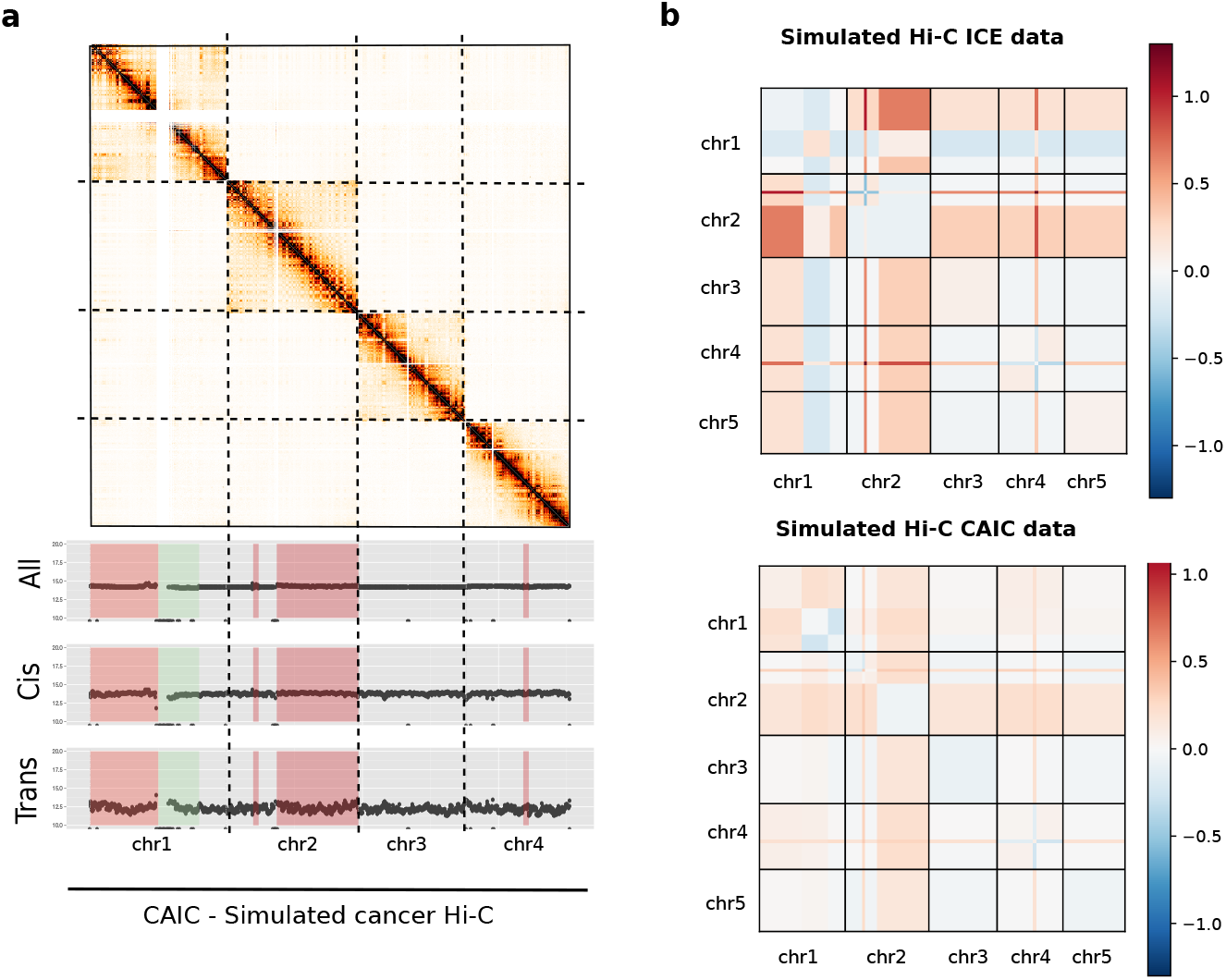
CNV-adjusted normalization of cancer Hi-C data. **a.** Hi-C contact maps of the four first chromosomes of our highly rearranged simulated data, together with the 1D signal of all, cis and trans data. Regions in red and green correspond to simulated gain and loss. **b.** Block-average error matrix of simulated ICE and CAIC Hi-C data. The CAIC efficiently removed the CNV effect, whereas the ICE normalization does not allow to correct for its effect.

Altogether, those results demonstrate the CAIC normalization procedure effectively removes copy-number effect.

### Application to breast cancer Hi-C data

A number of studies performed Hi-C experiments on cancer samples or cell lines [Barutcu et al., 2015, Taberlay et al., 2016, Le Dily et al., 2014]. We further explore the effect of our normalization procedures on two previously published Hi-C data from breast cancer cell lines: T47D [Le Dily et al., 2014] and MCF7 [Barutcu et al., 2015]. We process the T47D and MCF7 samples from raw data files to raw contact maps using the HiC-Pro pipeline [Servant et al., 2015]. As already seen in our simulation data (Figure 2a), we observe a strong copy number effect on the raw contact maps with respectively higher/lower contact frequency on gained/lost DNA regions in the T47D and MCF7 samples (Figure 5, Figure S9). Applying ICE on these data set does not remove entirely the copy number effect, and tends to flip the coverage profile between gained/lost regions in *cis*, therefore validating our previous observations on simulated data.

**Figure 5:**
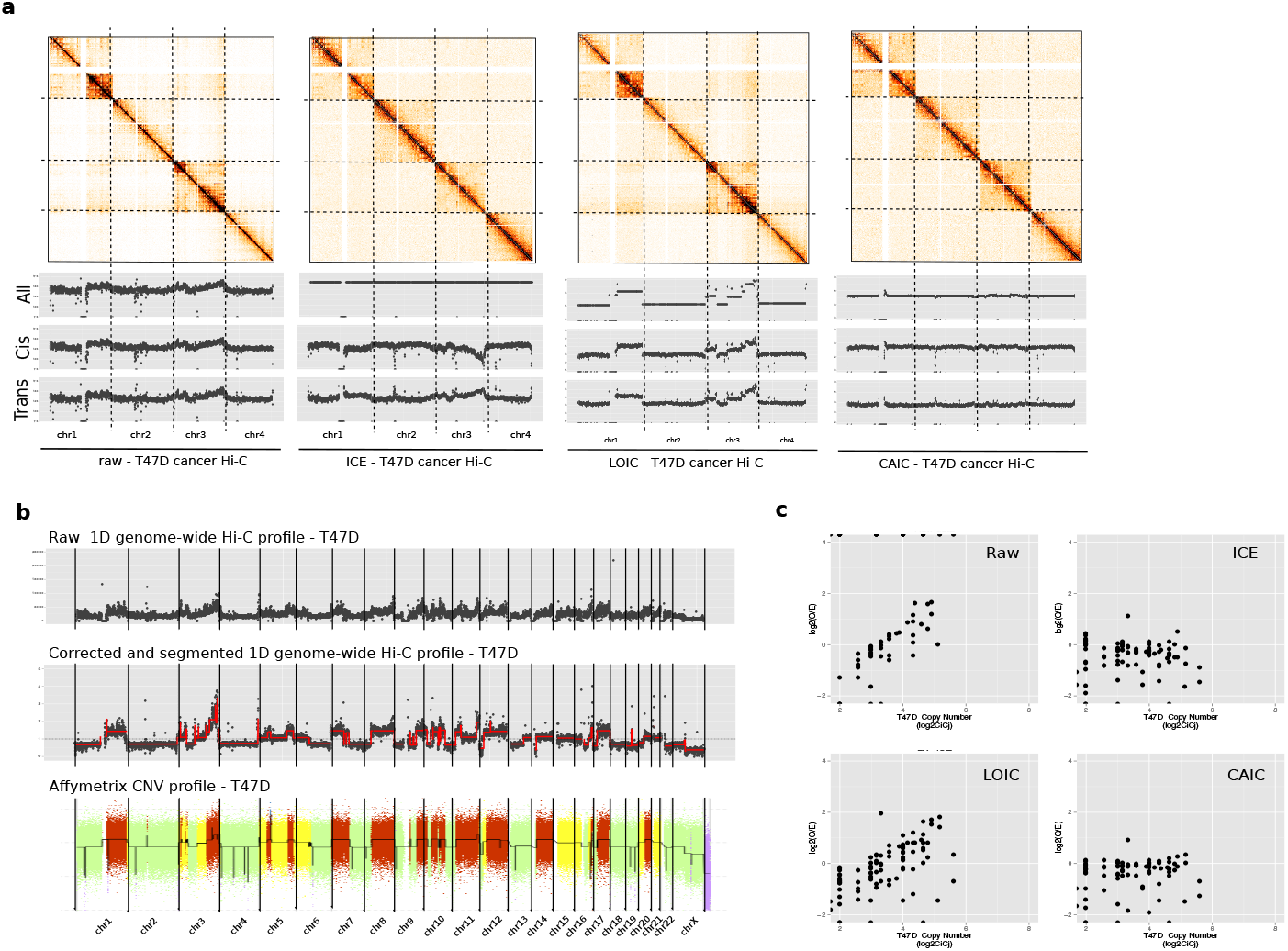
Normalization of T47D Hi-C data. **a**. Hi-C contact maps (250 Kb resolution) of the first four chromosomes of T47D cancer Hi-C sample. When looking at the 1D *cis* and *trans* profiles, we observed that ICE introduces a bias in the normalized data, therefore validating the observation made on the simulated data. We then applied the LOIC and CAIC normalizations in order to efficiently correct the data from systematic bias, while removing or keeping the CNVs effect. **b**. Estimate the copy number signal from the Hi-C data after correction and segmentation of the 1D profile. The inferred copy number signal from the Hi-C data are highly correlated with the copy number profile from Affymetrix SNP6.0 array. **c**. Correlation of raw and normalized contact frequencies with the copy number.

In order to estimate the copy number signal from these cell lines, we segment the 1D Hi-C profile as previously described. Interestingly, on both T47D and MCF7 data we observe a very good correlation between the copy number signal extracted directly from the Hi-C data and the copy number profile extracted from SNP6 Affymetrix array (Figure 5b, Figure S9b, Spearman cor=0.87 for both MCF7 and T47D data) We then apply the LOIC strategy as presented above so that the sum of each column/row follows the segmentation profile extracted from the data. As expected, we observe that the LOIC normalized contact maps conserves the copy number properties, and that the biases introduced by the ICE normalization no longer hold true. In addition, we also applied the CAIC normalization to correct of the copy number signal. Looking at the correlation between Hi-C counts and the copy number signal validates the efficiency of the methods (Figure 5c, Figure S10). In conclusion, applying the LOIC and CAIC methods on both cancer data set allows us to correct for systematic bias while conserving or removing the copy number structure.

### Normalization of capture-Hi-C data with genomic duplication

In addition to cancer data, we also investigate the relevance of LOIC to normalize Hi-C data in the presence of local structural events. Recently, Franke et al. [2016] investigated the effect of local duplications on chromatin structure and, in particular, on the formation of new topological domain. We therefore process the capture Hi-C data of the Sox9 locus and generate the raw and ICE contact maps at 10kb resolution. We focuse our analysis on samples with inter-TAD (dup-S) and intra-TAD (dup-L) duplications. In the context of inter-TAD duplication, Franke et al. [2016] described the formation of a new domain in the duplicated region by comparing the raw contact maps of wild type (WT) and dup-L samples (Figure 6c-d). Interestingly, when we compare the WT sample and the samples with the duplication events normalized by the ICE method, we observe that the ICE normalized maps do not allow duplication effects to be observed clearly. This is in agreement with our previous observations on cancer Hi-C data. We then apply our LOIC strategy following the observed 1D coverage profiles. As illustrated in Figure 6c, the LOIC normalization allows to remove systematic biases from the contact maps while keeping the copy number effect. Normalized maps allow to clearly observe the effect of both intra and inter-TAD duplication, validating the interest of the method to study local structural rearrangements.

**Figure 6:**
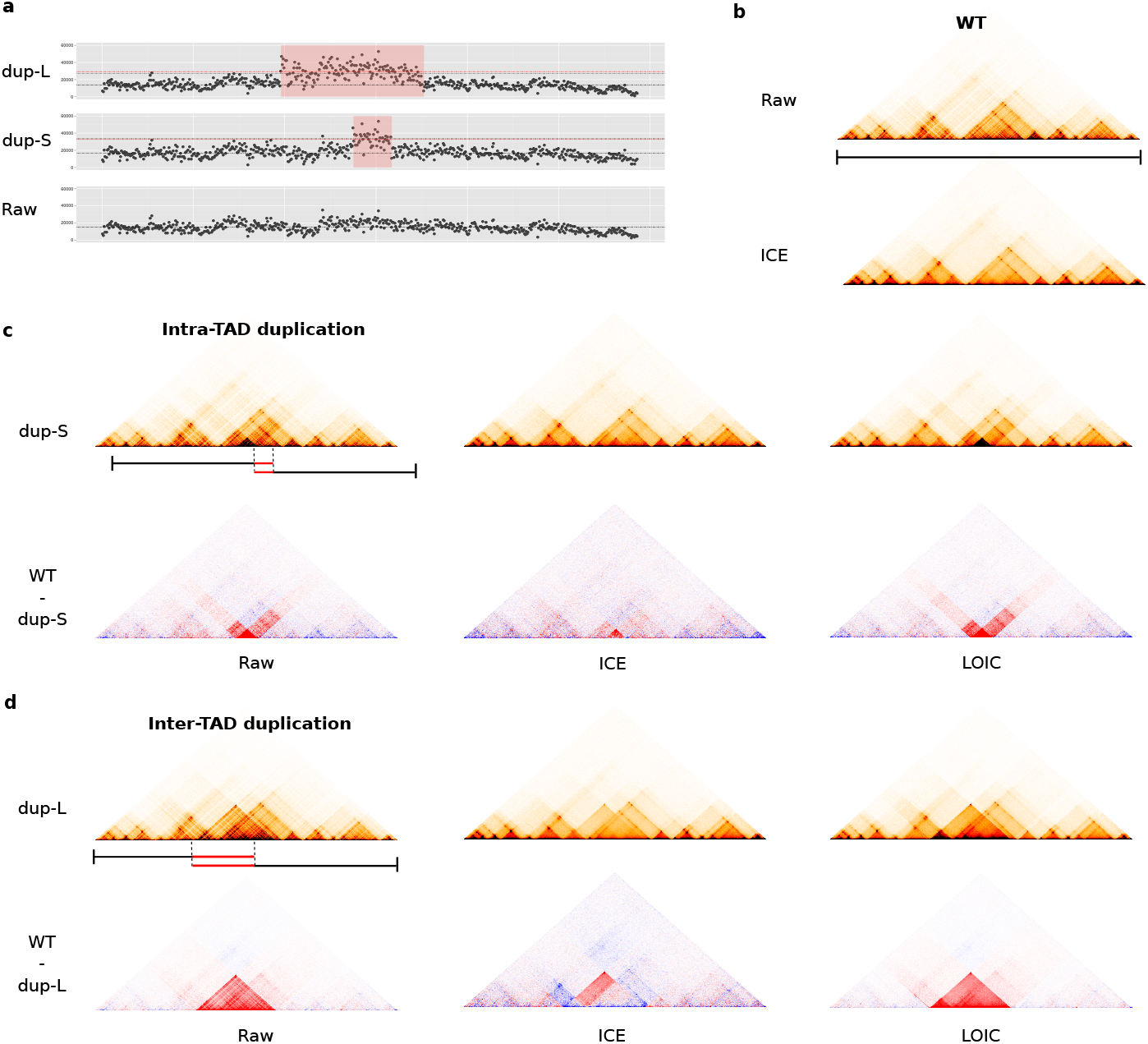
Duplication in capture-Hi-C and normalization. **a**. 1D profiles of capture-Hi-C wild-type sample (WT), with intra-TAD duplication (dup-S) or with inter-TAD (dup-L) duplication Franke et al.. As expected, the duplication samples are characterized by twice more contacts at the duplicated sites. **b**. Raw and ICE normalized contact maps of WT sample. c. Normalization of the dup-S sample with the ICE and LOICE methods. The duplication effect is visualized by subtracting the normalized WT and dup-S contact maps. **c**. Same approach applied to the dup-L sample.

### Removing the CNVs signal avoids misinterpretation of the chromosome compartment calling of cancer Hi-C data

We then further explore the impact of CNVs and normalization on the chromosome compartment calling. Looking at intra-chromosomal contact maps, the chromosome compartment profile appears as a checker-board-like interaction pattern, shifting from blocks with either high and low interaction frequency. Thus, chromosome compartments are usually detected using a Principal Component Analysis (PCA) on the correlation matrix of the distance-corrected intra-chromosomal contact maps. The first principal component then distinguishes the active (A) from inactive (B) compartments [Lieberman-Aiden et al., 2009] (see Supp Methods).

We perform this compartment calling analysis on the MCF7 Hi-C data normalized by ICE, LOIC, or CAIC methods and integrate the results with the histone marks data obtained from the ENCODE project [Dunham et al., 2012]. Surprisingly, we observe that, for most chromosomes, the compartment calling is not affected by the CNVs (Figure S12, Figure S13, Figure S14).

We then assess how the A/B compartments correlate with the active and repressive histone marks genome-wide (see Supp Methods). We observe that active compartments are associated with open-chromatin marks such as H3K27ac, H3K36me3 and H3K4me, and this, whatever the normalization method used. Respectively, inactive compartments are associated with repressive marks such as H3K27me3 or H3K9me3 (Figure 7a). In addition, we observed that the A-type and B-type compartments are very similar regardless the normalization used.

**Figure 7:**
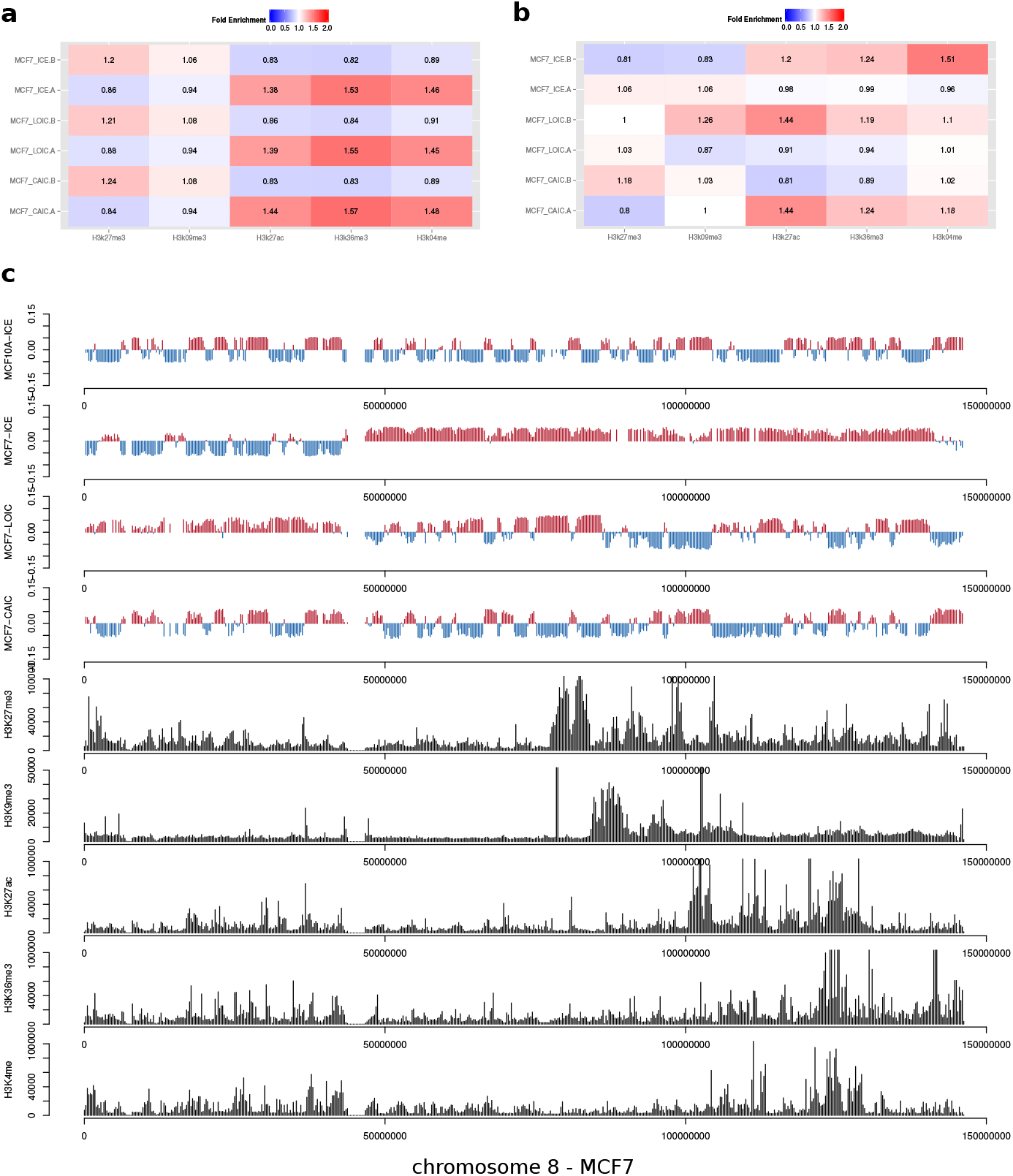
Detection of chromosome compartments. **a** Genome-wide enrichment of ChIP-seq histone marks in active (A) or inactive (B) compartments. Active compartments are enriched in open-chromatin marks, whereas inactive compartments are enriched in repressive marks. The results are concordant genome-wide, whatever the normalization method applied. **b**. Histone marks enrichment on chromosome 8 of MCF7 sample. On this chromosome, the copy number has a strong impact on the compartment calling. **c**. Results of the compartment calling for the chromosome 8 of MCF7 sample (first principal component), together with normalized ChIP-seq tracks. Active domains are in red. Inactive domains in blue.

However, looking at each chromosome independantly shows that, in the MCF7 data, the chromosome 8 harbours distinct compartment patterns according to the normalization method used. In this case, it is clear that the copy number affects the PCA analysis and the compartment calling (Figure 7b,c). We therefore conclude that it is important to correct for the copy number effect before running such analysis. Applying the CAIC normalization method outperforms the other methods, allowing to efficiently detect the A/B compartments on chromosome 8. The compartments pattern of the chromosome 8 extracted from the ICE normalized data is concordant with our previous conclusion that ICE is not appropriate to correct for CNVs, potentially leading to a wrong interpretation of the compartment profile. However, we also notice that, in this case, the A/B compartments can be rescued by looking at the second principal component of the PCA.

Altogether, these results demonstrate that, although the compartment calling seems globally not affected by the copy number effects, applying the CAIC strategy to normalize the data improves the detection of A/B compartments and avoids potential issues in their interpretation.

## Discussion

Chromosome conformation techniques are promising exploration tools to investigate links between the three-dimensional organization of the genome and functional and phenotypical effects in diseases. However, in the context of genomic rearrangements, dedicated methods should be applied as a novel source of signal related to copy number can arise. Predicting the exact effect of copy number variations on chromosome structure is a challenging task, as many factors influence the 3D structure of the genome and the resulting contact frequencies.

In this paper, we propose a simulation model to explore the effect of large copy number on Hi-C data. Starting from a diploid data set, our model is able to predict the effect of large copy number changes on interaction patterns. Although it represents a powerful tool to assess the ability of a method to deal with the copy number changes, it also demonstrates that predicting the real effect of copy number changes is very challenging. As an example, our model considers the duplicated regions as non-tandemly rearranged events, which, in some cases, certainly underestimates the intra-chromosomal effect of copy number. In addition, it does not integrate any biological knowledge and is therefore not designed to simulate changes due to the alteration of regulatory elements such as insulator regions. We also note that the sub-sampling strategy that we applied requires a high resolution diploid Hi-C data set in input. However, as the Hi-C sequencing depth is still increasing, this should no longer be a limitation.

Using our simulation model, we then demonstrate that applying ICE to data sets with abnormal karyotypes leads to unbalance corrections between amplified and lost regions. We further validate this observation using two Hi-C breast cancer data set.

We therefore propose two new normalization methods, respectively able to conserve or to remove the copy number, while correcting the data from other systematic biases. In addition, we also propose a segmentation procedure to directly extract the copy number signal and the breakpoints from the Hi-C contact probabilities and to use this information in the normalization process. Although this step is crucial for the normalization and can be challenging for noisy samples, our procedure performs well on all the data set used in this study, including the normal diploid samples.

The choice of the normalization method mainly depends on the context and biological questions. As an example, we demonstrate than, although the chromosome compartments calling and the PCA analysis are not dramatically affected by the copy number changes on MCF7 data, it can also lead to wrong interpretation if the data are not properly normalized. From our experience, this effect can be more important on other cancer types and mainly depends on the copy number profile of the tumor (data not shown). We therefore demonstrate the interest if our CAIC normalization method to remove the copy number biases, and to improve the reliability of the compartment calling.

In addition, we also illustrate the interest of keeping the CNVs information when looking at local structural rearrangements. In this context, we demonstrate that our LOIC method can easily be applied to properly remove the other systematic biases while keeping the copy number structure.

Taken together, our analysis highlights the importance of using dedicated methods for the analysis of Hi-C cancer data. It therefore paves the way to further explorations of the three-dimensional architecture of cancer genomes.

## Methods

Let us first introduce some notations. Given a segmentation of the genome into *n* genomic windows (or bins), Hi-C data can be summarized by a *n*-by-*n* symmetric matrix *C*, in which each row and column corresponds to a specific genomic loci and each entry *C_ij_* the number of times loci *i* and *j* have been observed in contact. Let *K ∈* ℝ^*n*^ the copy number profile of the sample of interest, which we represent as a piecewise constant vector.

We denote by *s*(*i, j*) the genomic distance between the loci, defined as the number of base pairs between the center of the two loci; if *i* and *j* are not part of the same chromosome, we extend this definition by setting *s*(*i, j*) = ∞ In this paper, we derive different ways to normalize the raw count matrix *C*: we denote by *N ^y^* the contact count matrix normalized with method *y* (*e.g. N* ^ICE^ represents the ICE normalized count matrix).

### Simulation of cancer Hi-C data

Before we turn to how to appropriately model cancer Hi-C data, let us first review some terminology. In the literature, *cis*-contact counts refers to the contact counts between two loci of the same chromosomes: this includes intra-chromosomal contact counts but also inter-chromosomal contact counts of homologous chromosomes. In this paper, we restrict the use of *cis*-contact counts to contact counts issued from the same DNA fragment, and we denote by “*trans*-homologous” (*transH*) interactions, the interactions between homologous chromosomes. Note that *cis* and *transH* contact counts are mostly indistinguishable (with the exception of allele-specific Hi-C) in Hi-C data hence the simplification of terminology usually used.

We now return to the problem at hand: how to simulate a contact count matrix *C*^sim^ of a cancer genome with abnormal copy number from a raw diploid contact count matrix *C*. In order to model the change in contact count abundances due to copy number variation, we first need to understand precisely which interactions are observed in the case of a simple diploid genome. For that purpose, we denote by *E_ij_* the expected contact count between loci *i* and *j*, and 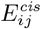, 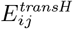 and 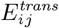 the expected *cis*, *trans* and *transH* -contact counts between *i* and *j*.

- if loci *i* and *j* belong to the same chromosome, the expected contact count *E_ij_* is the sum of (1) *cis*-counts from either of the homologous chromosomes; (2) the *transH* -counts between the two homologous chromosomes:

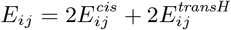
- if loci *i* and *j* belong to different chromosomes, then the observed contact counts *E_ij_* is the sum of either of the four possible *trans* interactions:

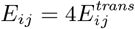 This can be generalized to polyploid genome or to the context of chromosomal abnormalities (Figure S1).
- if loci *i* and *j* belong to the same chromosome, let *k* be the number of *cis* interactions. If *i* and *j* belong to the same DNA segment, *k* = *K_i_* = *K_j_*. When *i* and *j* belong to different DNA segments, *k* could in theory take values between [0 *− min*(*K_i_, K_j_*)]. Here, we simulated the data with *k* = 2, or *k* = *min*(*K_i_, K_j_*) if *K_i_* < 2 or *K_j_ <* 2. Then

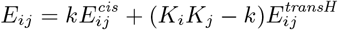
- if loci *i* and *j* belong to different chromosomes, then

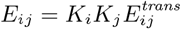

Now that we have derived how contact counts are decomposed in terms of *cis*, *transH* and *trans* contact counts, we can leverage those relationships to simulate the effect of copy number variations on contact count matrices.

In order to derive a scaling factor *p_ij_* which incorporates the copy number effect, we need to estimate 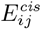, 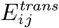and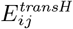, which is impossible without further assumptions. In practice, little is known about the probability of contact between homologous chromosomes, which is therefore difficult to estimate. However, we know that the chromosomes usually occupy their own space (chromosome territories) within the nucleus. We therefore make the assumption that all chromosomes are independent, and that the contact probability between homologous chromosomes can be estimated using the *trans* interaction between non-homologous chromosomes. We thus consider that 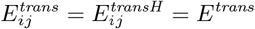

From these relationships, we then calculate the scaling factor *p_ij_* as following (recall that *E_ij_* is the expected copy number between *i* and *j* on genome with abnormal chromosomal interactions):

- we estimate *E^trans^* as the median *trans*-contact count;
- 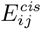 = *C_ij_* − *E^trans^*;
- Eij is estimated using the equations derived above;
- if loci *i* and *j* belong to the same chromosome, 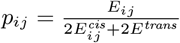
- if loci *i* and *j* belong to different chromosomes, 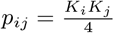

We thus obtain, for each entry of the contact count matrix *C_ij_*, a ratio *p_ij_* corresponding to the expected factor of enrichment or depletion of interactions for the loci *i* and *j*. In order to make the estimation of *p_ij_* more robust, we estimate it constant per blocks of identical copy numbers by taking the median of the empirical values in each block. (See Figure S11). Thus, the factor matrix *p* can be assumed to be block constant between regions of identical copy number variations. We thereby smooth *p* by computing the median scaling factor of block of similar copy number.

Finally, the simulated contact counts 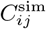 are generated by a binomial subsampling strategy of *C_ij_* by a probability equal to 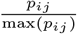 [Wiuf and Stumpf, 2006]:

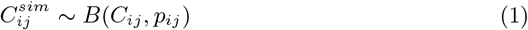

The reason for choosing a binomial subsampling as opposed to a simpler multiplication of the original Hi-C counts by a CNV-dependent factor, is that if *C_ij_* follows a Poisson or Negative Binomial distribution, then 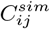follows the same distribution with modified expectation [C.Wiuf and P.H.Stumpf]. One limitation of this model is that the simulated counts can only be smaller than the original counts, which may be problematic if we start from small counts. It thus requires a diploid Hi-C data set with a sufficient sequencing depth to apply the downsampling strategy.

### LOIC: Correcting technical biases of Hi-C cancer data

To normalize the contact count matrix, we adapt the ICE method proposed by Imakaev et al. [2012] to incorporate the copy number effect. In particular, we use similar assumptions. First, the bias between two regions *i* and *j* can be decomposed as the product of two region-specific biases *β_ij_* = *β_i_β_j_*.

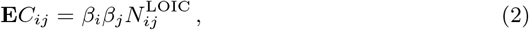

where *β ∈* ℝ^*n*^ is a vector of bin-specific *biases*, such as gc-content, fragment lengths, mappability, etc.

Second, all copy-number identical regions interact as much: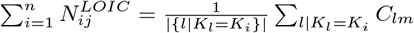 We refer to the second hypothesis as the “local equal-visibility assumption” to contrast it with ICE’s “equal-visibility assumption”: instead of enforcing an interaction profile constant across all the genome, we enforce an interaction profile constant for regions of identical copy number number.

Similarly to Imakaev et al. [2012], this problem can be solved exactly using matrix-balancing algorithms (under the assumption that the matrix is full decomposable [Sinkhorn and Knopp, 1967]). Note that if there is no copy number variations, this boils down to solving exactly the same problem as ICE. On the other hand, in the presence of copy number variation, the resulting interaction profile will be a constant piecewise function, whose value depends on the copy number of the two loci.

In order to apply the proposed method, one needs to know *a priori* the set of bins with a given copy number or the copy number breakpoints. It can either be found via probing the samples to estimate it using specific technologies or through prior knowledge on the cell-line or sample studied. When none of these options are available, we can leverage the information provided by Hi-C data directly to estimate it.

### Estimation of copy number from the contact count matrix

The copy number signal can be directly inferred from the Hi-C data in two steps. We first calculate the one-dimensional (1D) signal as the sum of genome-wide contact per bin, assuming that this signal reflects the true contact frequencies including the systematic Hi-C biases and the CNVs signal. We further calculate the GC content, the mappability and the effective fragment length of each bin end as already proposed [Hu et al., 2012]. The local genomic features of all chromosome bins are defined as the average of the corresponding features among all overlapping fragment ends. We then apply a Poisson regression model to correct the signal from GC content, mappability and fragment length, using the model proposed by Hu et al. [Hu et al., 2012]. The corrected profile is obtained by subtracting the fitted values to the observed data, and rescaled to be centered on 1. The normalized 1D data are then segmented using a pruned dynamic programming algorithm [Picard et al., 2011]. The segmented profile is smoothed with the GLAD package [Hupé et al., 2011] in order to optimize the breakpoint locations and to remove false positives events. The segmentation is an important step of the method which may need to be adjusted according to the signal-to-noise ratio of the data. In this study, we apply the same parameters to all data sets and we consider the smoothed line after the segmentation as the Hi-C derived copy number profile.

### CAIC: Removing the copy number effect

The previous section describes how to normalize the raw contact counts matrix *C* to adjust for unwanted variations such as GC-content, mappability, fragment lengths, while keeping the copy number information. We now propose to estimate the effect of copy number variations on the contact count matrix to offer the possibility of removing it. We denote by *N ^CAIC^* the normalized contact count matrix where the CNV effect has been removed.

We assume that the copy number effect for each pair of loci is identical for element with identical copy-number variations. This reflects that the copy-number effect between loci *i* and *j* is related to the amount of genetic material of those two regions, and thus identical between all pairs with similar copy-number variations. We can thus model the normalized contact count matrix as the product of a block-constant matrix *B* and corrected matrix 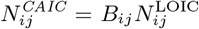, where each block is a function of the copy number in *i* and in *j*.

In addition, we assume that, on average, each pair of loci interacts roughly the same way as any pair of loci at the same genomic distance *s*:

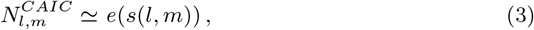

where *e*(*s*) is the expected contact count at genomic distance *s*. We leverage this assumption to cast an optimization problem:

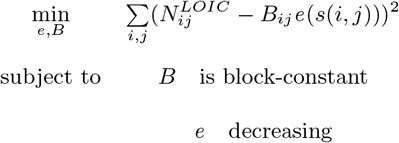

We solve this optimization problem genome-wide by iteratively estimating the block constant matrix *B* and the expected counts function *e* using an isotonic regression. Note that the *trans* estimation can be done jointly on the whole genome independently from the *cis* estimation.

## Availability of supporting data and code

All codes and data are available at https://github.com/nservant/cancer-hic-norm

## Authors’ contributions

NS, NV, EH, JV and EB designed the project. NS, NV and JV designed the simulation model, and the concepts of the normalization methods. NV implemented the normalization approaches. NS performed Hi-C data analysis. All authors read and approved the final manuscript.

## Acknowledgments

We would like to thank D. Gentien and the Institut Curie genomic platform for providing the MCF7 and T47D Affymetrix data. Many thanks to P. Gestraud, P. Hupe, F. Picard, G. Rigaill for their help in defining the segmentation strategy, and to J. van Bemmel for her feedbacks about the manuscript.

